# Improvements in upper extremity function in children with unilateral spastic cerebral palsy after intensive training correlates with interhemispheric connectivity

**DOI:** 10.1101/609313

**Authors:** Maxime T. Robert, Jennifer Gutterman, Claudio L. Ferre, Karen Chin, Marina B. Brandao, Andrew M. Gordon, Kathleen Friel

**Affiliations:** Burke Neurological Institute, Weill Cornell Medicine, New York, United States; Department of Biobehavioral Sciences, Teachers College, New York, United States; Graduate Program in Rehabilitation Sciences, School of Physical Education, Physical Therapy and Occupational Therapy, Universidade Federal de Minas Gerais, Belo Horizonte, Minas Gerais, Brazil

**Keywords:** Corpus Callosum, cerebral palsy, intensive interventions, Diffusion MRI, interhemispheric connections

## Abstract

**AIMS:** The corpus callosum (CC) regulates signalling between the two hemispheres and plays an important role in upper limb functions. There is limited evidence on the relationships between the integrity of the CC and upper limb functions in children with USCP. Furthermore, the extent of how much the CC can be used as a biomarker to predict hand functions following intensive interventions remains unknown. We examined 1) the relationship between hand function and tractography of the CC, and 2) the associations between the integrity of the CC and changes in hand function following intensive intervention.

**METHODS:** Forty-four participants received 90 hours of intensive therapy and were randomly allocated in one of two training groups: Hand-arm Bimanual Intensive Therapy (HABIT) or Constraint-Induced Movement Therapy (CIMT). Hand functions were assessed pre-and post-intervention by a blinded clinician using the Jebsen-Taylor of Hand Function (JTTHF), Assisting Hand Assessment (AHA), and Box and Blocks test (BBT). Functional goals and daily functioning were measured using the Canadian Occupational Performance Measure and the Abilhand-Kids. CC tractography was reconstructed using diffusion tensor imaging (DTI). Corpus callosum was segmented into three regions of interest (genu, midbody and splenium). Linear regression and pearson correlations were used to assess the relationships between bimanual outcomes and DTI parameters.

**RESULTS:** Both groups demonstrated improvement of hand function (p<0.05). JTTHF, AHA and BBT significant correlated with DTI variables for all ROIs (p<0.05). Bimanual and perceived manual ability of children changes following CIMT were negatively correlated with number of streamlines and number of voxel for the whole CC (r=-.442, p=0.05), midbody (r=-.458, p=0.042) and spelnium (r=-.512, p=0.021). No significant correlation was observed for the HABIT group.

**INTERPRETATION:** Tractography of the CC was found to be associated with unimanual and bimanual functions at baseline. Children with reduced integrity of the CC and with greater bimanual impairments improve more from CIMT. On the contrary, all children in the HABIT group had similar improvements independent of the CC integrity.

## Introduction

Many activities of daily living (ADL) require the use of both hands. However, in children with unilateral spastic cerebral palsy USCP, these activities could be limited due to structural abnormalities of the brain and a wide range of sensorimotor impairments including weakness in one arm, reduced bilateral coordination and reduced sensation [1, 2]. Several interventions have been proposed with the ultimate goal to increase their functional independence [3].

Among these, constraint induced movement therapy (CIMT) and hand-arm bimanual intensive therapy (HABIT) have been widely accepted to be effective interventions to increase hand function [3]. While both therapies have a similar objective, the approach is different as CIMT focuses on unimanual dexterity, whereas HABIT focuses on increasing the capacity of bimanual performance [4, 5]. Despite their general efficacy, a high level of variability in hand improvements is often observed. To elucidate this, recent studies have looked at possible biomarkers that could explain this observed variability in hand functions following intensive intervention.

Possible biomarkers to predict hand functions include sensory and motor connectivity due to their role in skilled movements [6, 7]. Recent studies using advanced neuroimaging techniques, such as diffusion tensor imaging, indicated that the integrity of the sensorimotor connectivity is associated with the severity of hand function impairments in children with USCP [8–13] These results suggest that upper limb function may depend on the integrity of these pathways. Only a handful of studies have partially supported that higher white matter integrity of the CST is associated with positive changes of hand functions following intensive intervention [14].

Furthermore, the efficacy of an intervention on hand function may also depend on the lateralization of the corticospinal tract (CST) [15] although the latter was found not to be a predictor of upper limb functions following HABIT [6] or CIMT [16]. Nonetheless, the result of these studies reinforces the importance to study other possible structures involved in hand function changes [8, 13, 17].

The corpus callosum (CC), which is the primary structure that integrates signaling from both hemispheres, may be one of the possible biomarkers for bimanual skilled movements [8]. Children with USCP have clear structural abnormalities of the corpus callosum (i.e., reduced number of streamlines and fractional anisotropy), which may explain the poor bimanual performance [8, 18, 19]. Among possible factors that may affect hand performance, ipsilesional and contralesional hand dexterity has been associated with the motor area of the CC [10]. Moreover, higher white matter integrity (higher fractional anisotropy; FA) of the genu and midbody was correlated with better manual function [8, 17]. Yet, the relationship between the integrity of the corpus callosum and changes in bimanual function following intensive treatment remains largely unexplored.

Thus, the purpose of the present study was to 1) assess the relationship between the integrity of the white matter of the CC and hand function, 2) to relate the clinical changes to the integrity of the white matter of the CC. Our exploratory objective will assess changes of the CC following the intensive intervention. Our first hypothesis is that the integrity of the CC will be positively correlated with unimanual and bimanual function. Our second hypothesis is that integrity of the CC will be positively correlated unimanual function and bimanual function following CIMT and HABIT, respectively. For our exploratory objective, we hypothesized that FA values and number of streamlines of the midbody will increase following both CIMT and HABIT.

## Methods

### Participants

A total of 44 children (24 males, 20 females, age 6 to 17) were randomized into one of the two intensive interventions: Hand-Arm Bimanual Intensive Therapy or Constraint-Induced Movement Therapy. Participants were recruited from our Web site (http://www.tc.edu/centers/cit/) and online communities. All participants were invited to receive an on-site physical examination or an examination videotaped by their caregiver or physical or occupational therapist. The inclusion criteria were: 1) age 6 to 17 years, 2) diagnosed with unilateral spastic cerebral palsy 3) capable of participating in 6 hour long day camp while separated from parent, 4) capable of following directions regarding hand use and testing, 5) capable of communicating needs, 6) able to lift the more-affected arm 15cm above a table surface and grasp light objects. Exclusion criteria included: 1) unwillingness to comply with instructions or other behavioral issues making delivery of an intensive therapy infeasible, 2) health problems unassociated with hemiplegia, 3) visual impairment that could interfere with participation, 4) orthopedic surgery on the more-affected hand within 1 year, and 5) seizure disorder, 6) botulinum toxin in the more affected upper extremity within the past 6 months or intended treatment within the study period. Informed assent and consent forms were obtained from participants and caregivers, respectively. This study was approved by the Institutional Review Boards of Teachers College, Columbia University and Burke Neurological Institute, Weill Cornell Medical College.

#### Procedures

##### General Procedures

A crossover design was used where assessments were collected before the Hand-Arm Bimanual Intensive Therapy (HABIT) and the Constraint-Induced Movement Therapy (CIMT) training to establish a baseline, immediately and 6 months after the intervention. A total of five intensive training day-camps were conducted at Columbia University from July 2013 to August 2017. Each camp included 8 to17 children with USCP. Participants were engaged in treatment 6h/d for 15 consecutive weekdays (90 hours) during school recess by trained interventionists. These included physical and occupational therapists, graduate students in kinesiology/neuroscience, speech pathology, and psychology, and undergraduates. The interventionist training, administered by the supervisors, was standardized during a 2-hour session based on established manual of procedures. Camp training focused on strategies to engage children in use of hands, safety and data logging procedures. Additional ongoing training was provided during the interventions and daily team meetings. The camp room had supervisors (including a physical and occupational therapists experienced with the treatment), responsible for ensuring treatment fidelity. The supervisors modeled and ensured uniformity of treatment. Room design permitted participants to work individually with their interventionist or in groups (1:1 interventionist:participant ratio was always maintained). Interventionists were paired with children based on age, gender, and caregiver input. Interventionists avoided verbal prodding to use the more-affected hand, and instead provided tasks necessitating the use of both hands and established rules prior to each activity. This allowed the child to choose which hand to use for different components of a bimanual activity. Children were engaged in fine and gross motor activities individually chosen according to the child’s abilities, impairments and improvements deemed possible by the supervisors in order to achieve success and to encourage bimanual use, as well as child’s active problem solving. Task difficulty was progressively graded focusing on increasing the level of difficulty, speed, or accuracy according to a child’s improvements, or spatial and temporal constraints of the activities. The procedures included the use of whole task practice (sequencing successive movements in the context of activities, such as self-care or play activities), part task practice (practice of specific components of the task in a repetitive sequence of 30 seconds) and individualized functional goal training. The activities chosen for goal training were selected according to the parents’ priorities, reported using the Canadian Occupational Performance Measure (COPM) at the pretest assessment. For additional details about HABIT and CIMT, see [5] and [4, 20].

#### Behavioral Measures

Participants were evaluated directly prior to treatment (pre), within 2 days (post) and 6 months after treatment by a physical therapist blinded to the training. Five outcome measures were used to quantify unimanual capacity, bimanual performance and functional goals.

Unimanual dexterity was assessed using two tests: (1) the Jebsen-Taylor Test of Hand Function (JTTHF) and (2), the Box and Block Test. The JTTHF is a standardized timed-test of simulated functional tasks quantifying the time to complete a battery of unimanual activities [21, 22]. The activities include flipping index cards, object placement, simulated eating, stacking checkers, and manipulating empty and full cans. Reliability for children with non-progressive hand disabilities is high. The Box and Block Test measures the number of blocks moved between the two boxes in 60 seconds [23]. Both unimanual dexterity tests were performed on each hand separately.

The Assisting Hand Assessment (AHA version 5.0) quantifies the effectiveness with which a child with unilateral disability uses his or her affected (assisting) hand in bimanual activity [24, 25] and has excellent validity/reliability [26]. The test was videotaped and scored offsite by an experienced evaluator blind to the fact the children received HABIT. Data was reported in logit-based units (AHA-units). For the AHA, an improvement of 5 units is considered the smallest detectable difference [27].

The ABILHAND-Kids is a valid/reliable questionnaire assessing the perceived manual ability of children [28]. The test comprises a list of manual activities in which the caregivers score the amount of difficulty children may experience during their performance in activities of daily living that required hand use. Data was reported in logit-based units.

To establish/evaluate children’s functional goals, the COPM with caregivers [29] was conducted. The COPM identifies and measures changes in functional problems considered relevant by clients through interview, and is valid/reliable. The most relevant functional goals to be accomplished are defined, ranked in importance, and rated on performance and satisfaction. In this study, caregivers selected the goals and rated the child’s performance since these are abstract concepts for children of this age. For the COPM, a change of 2 points is considered clinically meaningful.

#### MRI Data Acquisition

Neuroimaging was performed on 44 participants. Due to excessive head movements (>5mm) during rest fMRI, we ask only twenty-four participants to come back after the intervention. The DTI was used to reconstruct the interhemispheric connections. MRI protocol was performed on a 3T Scanner (Siemens Magnetom Trio, Citigroup Biomedical Imaging Center, Weill Cornell Medical College). A total of 75 slices were acquired (matrix 112 × 112, FOV = 224 mm, 65 directions, b-value = 800 s/mm^2^, TR = 9000ms, TE = 83ms). The participants were positioned in a supine position with padding around the head to minimize the movement and reduce noise. The participants were not physically constrained nor received any sedation.

#### MRI Data Analysis

DTI analysis was performed using DTI Studio (John Hopkins University, Baltimore, MD, USA), which included fractional anisotropy, vector maps, and color-coded maps. An image was first created to mask the background noise at the threshold of 30 dB, using standard linear regression for tensor calculation. Images containing movement artefacts were excluded by visually inspecting the original images using the apparent diffusion constant function [30]. Reconstruction of the callosal pathways was done using the Continuous Tracking method [31]. Fiber tracking started < 0.15 and was terminated if tract turning angle was >70.

Regions of interest (ROI) was determined using anatomical location (XYZ) through orientation-based color-coding maps (red for fibers with medial-lateral orientation). The CC was segmented into three segments based on the Witelson parcellation scheme [32]. It was segmented into the genu (the anterior third), the midbody and the splenium (the posterior part) of the CC.

The following parameters were calculated: FA, number of steramlines and voxels. To investigate if noise correction could affect the results, the total of bad pixels for each slice at each gradient was first compared between pre and post assessment and then normalized with the total number of streamlines. Data was analyzed by a trained researcher (first author) and all of the data were re-analyzed by a different trained rater (second author). Also, data was re-analyzed by the same rater a month after the initial analysis for intra-rater reliability measurements.

##### Measures

###### Statistical Analysis

Statistical analysis was performed using SPSS (Version 25, Statistical Production and Service Solutions, Chicago, IL). Gaussian distribution was verified using a chi-square test. Reliability of the DTI measures was examined using the Cronbach Alpha coefficient. Multiple mixed linear models on test sessions were performed for all clinical outcomes with time as a fixed factorial factor [33] to see improvements over time. Mixed linear models allow the estimation of interindividual variability and intraindividual patterns of change over time while accounting for missing data. The time factor was the independent variable, while the dependent variables consisted of the JTTHF, the Box and Block Test, the AHA, the ABILHAND-Kids and the COPM. For the first objective, a series of Pearson correlations was conducted to study the relationship between clinical scores and DTI variables. For the second objective, correlations were conducted to investigate the relationship between changes in clinical outcomes and DTI variables for every ROIs. Multiple regression analyses were then conducted in the same model with the inclusion of the baseline AHA score as a predictor, which was fitted using the enter selection method. For our exploratory objective, several paired t-tests were performed to measure the DTI changes following the intensive intervention. Significance was set at p<0.05.

## RESULTS

### Patient Flow

A total of 65 individuals participated in the summer camps from July 2013 to August 2017. Of those, 44 participants were able to complete the baseline neuroimaging and only 20 of those were invited back at post due to excessive head movements and noise during the MRI at baseline.

There was a Gaussian distribution for all measures. Significant improvements in the clinical measures were observed from pre to post (p<0.05 for all measures). No significant differences were observed between the CIMT and HABIT groups at baseline for any clinical or DTI measures (p>0.05). All children completed the 90 hours of training.

### Diffusion Tensor Imaging Reliability

For the DTI measurements, both inter-reliability and intra-reliability were good to excellent using the intraclass coefficient ranging from 0.816 (CI 0.079-0.963) to 0.979 (CI 0.896-0.996) and from 0.746 (CI −0.267-0.949) to 0.988 (CI 0.940-0.998), respectively.

#### Objective 1

Data was combined as no differences were observed between the two groups at baseline (p<0.05).

### Manual dexterity

The JTTHF scores were significantly correlated with FA values, number of streamlines and number of voxels for all ROIs of the CC (p<0.05). The BBT scores were significantly correlated with number of streamlines and number of voxels for all ROIs of the CC (p<0.05). FA values of the genu were also significantly correlated with the BBT score (r=0.307, p=0.045).

### Bimanual

The AHA scores were significantly correlated with FA values (with the exception of the splenium), number of streamlines and number of voxels for all ROIs of the CC (p<0.05). *Functional goals and daily living*

No correlations with FA values were found (p>0.05). ABILHAND-Kids scores were correlated with number of streamlines of the CC (r=0.312, p=0.039) and the splenium (r=0.328, p=0.30). COPM score was correlated with the number of streamlines and number of voxels for all ROIs of the CC (p<0.05).

#### Objective 2

Three separate sets of analysis were performed for this objective such as the following 1) CIMT+HABIT, 2) CIMT and 3) HABIT.

#### CIMT and HABIT

Changes in ABILHAND-Kids were significantly correlated with the number of streamlines and voxels for the following ROI’s: CC, midbody and splenium (p<0.05). No other significant correlations were observed.

### Corpus Callosum

For changes in the ABILHAND-Kids, the multiple regression model with all two predictors explained 16% of the variance for the number of streamlines (F=3.893, p=0.028) and 15% for the number of voxels (F=3.634, p=0.035).

### Genu

The multiple regression model was not significant for the genu.

### Midbody

For changes in the ABILHAND-Kids, number of streamlines (F= 5.498, p=0.008) and number of voxels (F=4.908, p=0.012) explained 21% and 19% of the variance using the multiple regression model, respectively.

### Splenium

Number of streamlines (F=3.321, p=0.046) explained 14% of the variance using the multiple regression model for the changes in the ABILHAND-Kids.

#### CIMT

Bimanual and perceived manual ability of children changes were correlated with the number of fibers and voxels for the CC, midbody and splenium (p<0.05). No other significant correlation was observed (p>0.05).

### Corpus Callosum

For changes in bimanual function, FA values (F=6.265, p=0.009), number of streamlines (F=5.042, p=0.019) and number of voxels (F=5.089, p=0.019) explained 42%, 37% and 37% of the variance using the multiple regression model, respectively. Number of streamlines (F=5.056, p=0.019) and number of voxels (F=5.433, p=0.015) explained 37% and 39% of the variance using the multiple regression model for the changes in the ABILHAND-Kids.

### Genu

The multiple regression model did not show significance for the genu.

### MidBody

For changes in bimanual function, FA values (F=5.470, p=0.015), number of steramlines (F=5.186, p=0.017) and number of voxels (F=5.522, p=0.014) explained 39%, 38% and 39% of the variance, respectively. For changes in the ABILHAND-Kids, number of steramlines (F=4.667, p=0.024) and number of voxels (F=5.522, p=0.014) explained 35% and 34% of the variance, respectively.

### Splenium

For changes in bimanual function, FA values (F=5.338, p=0.0156), number of streamlines (F=5.237, p=0.017) and number of voxels (F=5.355, p=0.016) explained 39%, 38% and 39% of the variance, respectively. For changes in the ABILHAND-Kids, number of streamlines (F=6.933, p=0.006) and number of voxels (F=5.906, p=0.011) explained 45% and 41% of the variance, respectively.

#### HABIT

Changes in the COPM was correlated with the FA and number of steramlines of the genu (p<0.05). Change in bimanual function was correlated with the FA of the midbody (r=417, p=0.042). No other significant correlation was observed. No significant multiple regression model was observed for all ROIs.

#### Exploratory objective Objective 3

There was an increase of number of streamlines and number of voxels for the CC (p<0.001, p=0.001), midbody (p=0.023, p=0.040) and splenium (p=0.006, p=0.003). Noise correction was also similar between pre and post assessments (p>0.05).

##### Discussion and conclusion

The objectives of this study were 1) to characterize the relationship between the integrity of the CC and hand functions, and 2) to assess the relationship between changes in hand functions and baseline integrity of the CC. Our results supported previous findings by showing strong correlations between DTI variables and hand functions. Furthermore, children with poorer integrity of the CC were associated with better bimanual changes for CIMT, but not for HABIT.

Our results reinforced the notion that abnormal CC is associated with poorer unimanual and bimanual functions in children with USCP [8, 10, 17]. For example, while Weinstein et al. (2014) did not show any correlation between the integrity of the CC and unimanual function, a subsequent study with a larger sample size from the same authors indicated that higher FA values of the genu, midbody and splenium were associated with better performance on the JJTHF [17]. Bimanual functions measured by AHA were also found to be positively associated with lower mean diffusivity of the genu and midbody regions [17] and with number of streamlines of the splenium [8]. Surprisingly, FA values were associated with AHA scores, but only immediately and 6-week after the intervention. Interpretation of the results from Weinstein et al., (2015) paper is limited due to the small sample size, the heterogeneity of CP and the ceiling effect observed in four participants on the JTTHF. These factors may have eclipsed any possible significant correlation between the integrity of the CC and upper limb functions. Addressing these limitations, our results propose an important role of the CC in both unimanual and bimanual functions in children with USCP. While the exact mechanisms in the CC for unimanual and bimanual functions remain debatable, some noteworthy reviews have described the interaction between the two hemispheres and whether the regulation of the latter is done through inhibitory or excitatory signaling [34, 35]. Evidence of the involvement of the CC on unilateral and bimanual skills was unequivocally found in patients with lesions to the CC [36, 37], in individuals with stroke [38], callosotomized patients [39] and in animals studies [40]. Consistent with previous work, our results reinforce the importance of the CC in unimanual and bimanual functions [41–44].

In search of a biomarker to predict bimanual changes following intensive therapies, other sensorimotor pathways, such as the corticospinal tracts, have received considerable attention [6, 7, 15]. However, none of these studies has looked at the CC and its association with clinical changes following therapies. In children with USCP, interhemispheric connections appeared to be important for attention, memory and mood, but little is known in motor functions (and even less relating to the types of training). To the best of our knowledge, only one study evaluated the correlations between DTI measures and changes in upper limb functions following bimanual therapy [14]. Similar to our findings, no significant correlation was observed between bimanual changes and the integrity of the CC following bimanual therapy in children with CP. On the contrary, following CIMT, children with abnormal integrity of the CC had greater bimanual changes as opposed to those with higher integrity.

Two possible mechanisms may explain why CC could be a predictor of bimanual changes following CIMT, but not HABIT. First, CIMT may help to improve motor function by reducing the activation of the transcallosal inhibition pathway [45]. In this case, due to the poor integrity of the CC, interhemispheric inhibition is most likely minimal as compared to those with near-intact CC promoting better recovery. A previous study using inhibitory repetitive transcranial magnetic stimulation over the unaffected motor cortex in adults with stroke demonstrated improvement of motor function following CIMT [46]. The findings of this study reinforced the hypothesis that reduced activation of the inhibitory connection leads to better recovery. Looking at diffusion neuroimaging predictors of clinical improvement following a 3-week CIMT, one study in children with USCP also arrived at the same interpretation despite the lack of measure diffusion imaging of the CC and small sample size [7]. Secondly, the nature of activities practiced in HABIT reflects bimanual coordination unlike those practiced in CIMT. Thus, regardless of the integrity of CC, all children were able to improve bimanual functions based on the fact that more bimanual practice was done in the HABIT group. Together, these two explanations suggest that that integrity of the CC may only be used as biomarker following CIMT and not HABIT.

Following an intensive intervention, the number of streamlines and the number of voxels were measured in several regions of the CC. Our results need to be interpreted cautiously for several reasons. First of all, the exact mechanism of diffusion changes underlying neural micro-structures changes remains unknown. A recent review suggested that these changes may be due to several factors, including an increase in myelination, axonal packing and/or temporal development [47]. Comparison of studies using DTI measures may also be difficult as there is a variety of methods to analyze diffusion data and spatial resolution of the MRI may differ [47, 48]. At present, it is unclear what fiber counts actually measure. In fact, several factors, such as length, curvature and degree of branching of the pathways, undermine the direct interpretation of fiber count. Reporting the number of streamlines rather than the fiber count takes into consideration these factors as well as the tractography algorithm [48]. The estimated number of streamlines may be an oversimplified approach that may neglect potential micro-structural changes [48].

A sensitive period to promote neuroplasticity and an increase myelination in the CC has been suggested in young musicians under the age of 7 years old [49–51]. These studies are not discarding the possibility to have microstructural changes at a later age if the training is done through the optimal settings. The role of physical activity on the CC has only been studied in one study [52]. Using DTI, microstructural properties of the CC in 143 children aged between 7 to 9 years old was measured before and after a physical activity program. Children were randomly allocated into one of the two group: 9-month after-school physical activity program or to a controlled group. Increases in FA values and increased myelination measured with radial diffusivity were observed in the genu ROI for the physical activity program. Under the right circumstance, this study highlights the potential to have neuroplasticity throughout childhood. Nonetheless, our results suggest a possibility to see changes of the white matter structure in children with USCP. More robust studies with more sophisticated models as well as an increase of participants may shed the light on this interesting finding.

## Acknowledgments

The authors thank the children and the families who participated in this study. We also thank the volunteers in their assistance with data collection.

